# Topological linkage disequilibrium calculated from coalescent genealogies

**DOI:** 10.1101/286393

**Authors:** Johannes Wirtz, Martina Rauscher, Thomas Wiehe

**Affiliations:** Institut für Genetik, Universität zu Köln

**Keywords:** Linkage Disequilibrium, Topological Linkage Disequilibrium, Kingman Coalescent, Ancestral Recombination Graph, Sequential Markov Coalescent

## Abstract

We revisit the classical concept of two-locus linkage disequilibrium (*LD*) and introduce a novel way of looking at haplotypes. In contrast to defining haplotypes as allele combinations at two marker loci, we concentrate on the clustering of sampled chromosomes induced by their coalescent genealogy. The root of a binary coalescent trees defines two clusters of chromosomes. At two different loci this assignment may be different as a result of recombination. We show that the amount of shared chromosomes among clusters at two different loci, measured by the squared correlation, constitutes a natural measure of *LD*. We call this *topological LD* (*tLD*) since it is induced by the topology of the coalescent tree. We find that its rate of decay decreases more slowly with distance between loci than that of conventional *LD*. Furthermore, *tLD* has a smaller coefficient of variation, which should render it more accurate for any kind of mapping purposes than conventional *LD*. We conclude with a practical application to the LCT region in human populations.

## 1. Introduction

Maintenance and decay of haplotype structure and allelic association are of prominent interest when studying the evolutionary dynamics of recombining chromosomes. One possibility of quantifying this association is by the well-known concept of *Linkage Disequilibrium* (*LD*), reviewed in several articles, for instance [1, 2, 3]. *LD* measures the degree of statistical dependence among alleles at two - or more - loci and is commonly defined via a combination of haplotype frequencies. In the simplest non-trivial case two alleles at each of two loci can be distinguished. In the framework of the polarized infinite sites model [4], two allele classes - derived and ancestral - can be distinguished for each polymorphic site. The derived alleles can be traced back to a single mutation event, marking the origin of the derived class at this locus. This event may lie on any branch of the underlying coalescent tree [5] and divide the leaves in two classes: those belonging to the subtree rooted by the mutation, and those belonging to the complement. In this work, we consider the most recent common ancestor, i.e. the root of the entire coalescent tree, rather than of a single mutation event as such a class marker, which divides leaves into a ‘left’ (from the root) and a ‘right’ class. Along a chromosome tree topology may change as a consequence of recombination and thereby re-shuffle leaves among the two classes. This gives rise to an extended definition of linkage disequilibrium: considering two coalescent trees along a recombining chromosome, we call the correlation between left and right class members *topological linkage disequilibrium, tLD* for short. While *tLD* is a theoretically simple concept, coalescent topology is in practice generally unknown and has to be estimated from polymorphism data. Still, *tLD* has a number of interesting properties. For instance, *tLD* decays more slowly with distance than conventional *LD*, because only a subset of recombination events affects tree topology at the root. This is of practical interest when measuring long-range *LD* and searching for possible interactions between chromosomally distant loci. To estimate tree topology several SNPs - on the order of about ten or more - are required. However, this jointly considering of SNPs, rather than calculating *LD* for all SNP-pairs, introduces a smoothing effect compared to conventional *LD*. In fact, the coefficient of variation (CoV) of *tLD* is much smaller than that of *LD*, especially for short distances between loci. For larger distances, the CoV of *tLD* is almost constant and only slightly larger than 1, while the CoV of conventional *LD* is around 3, indicating a much higher dispersion relative to *tLD* (Figure 6).

## 2. Materials and Methods

### 2.1 Linkage Disequilibrium

Consider a population of constant size 2*N* chromosomes and let *α, β* be two loci with alleles *a, A* and *b, B* with allele frequencies *p*(*a*), *p*(*A*), *p*(*b*) and *p*(*B*). Let the four haplotypes *ab, aB, Ab* and *AB* have frequencies *p*(*ab*) = *x*_1_, *p*(*aB*) = *x*_2_, *p*(*Ab*) = *x*_3_ and *p*(*AB*) = *x*_4_. Two-locus linkage disequilibrium is

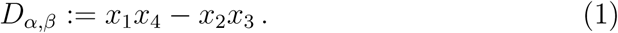

This can also be written as *D*_*α,β*_ = *x*_1_ -(*x*_1_ +*x*_2_)(*x*_1_ +*x*_3_) = *p*(*ab*) -*p*(*a*) *p*(*b*). A configuration of *x*_1_, …, *x*_4_ such that *D*_*α,β*_ = 0, is called *linkage equilibrium*. In this case, all haplotype frequencies are identical to the product of the involved allele frequencies. In practice, allele and haplotype frequencies in the population must usually be estimated from sample frequencies. In what follows, we do not distinguish between samples and populations and view all frequencies as population frequencies.

Since *LD* depends on allele frequencies, several standardizations have been introduced. One is Pearson’s correlation coefficient of the allelic association

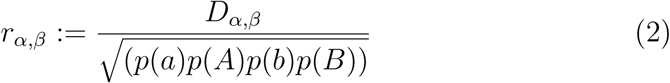

which can take values between −1 and 1, where the sign depends on labeling of alleles. Furthermore,

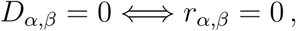

and

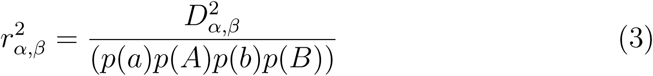

is the χ^2^-value of the allelic association. An alternative, frequently used, standardization is

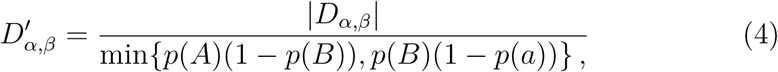

known as “Lewontin’s *D*” [2]. All three quantities, 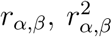 and 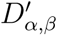 have been used to estimate recombination rates, divergence times between species, or to statistically test the neutral evolution hypothesis [6, 7, 8]. Here, we focus on 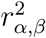.

Let *c* be the recombination probability per generation per chromosome between loci *α* and *β*. The random variable 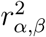 changes due to the action of drift and recombination.

We consider first its expectation with respect to *c*. When *α* = *β*, we have *x*_1_ = *p*(*a*) = *p*(*b*) and *x*_4_ = *p*(*A*) = *p*(*B*), and therefore

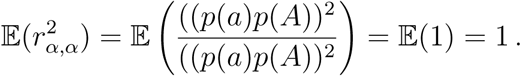

A fortiori, two identical loci do not recombine, i.e. *c* = 0. However, this may still hold also if *α ≠ β*, for instance in non-recombining chromosomes. In this case *LD* may take any value 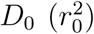 and, on average, remains at this value, regardless of time and initial allele frequency distribution, i.e.

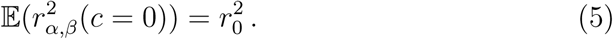

Let *x*_*i*_ (*D*) denote haplotype frequency (*LD*) in the current generation and 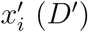 denote haplotype frequency (*LD*) in the next generation. We have

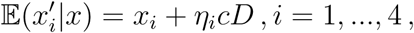

where *η*_1_ = *η*_4_ = *-*1 and *η*_2_ = *η*_3_ = +1, and therefore

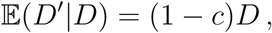

expected *LD* decreases at rate *c* per generation for any value *c >* 0. Since, on average, allele frequencies do not change across generations, we have

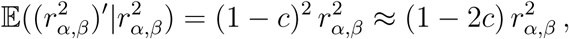

where the last approximation holds for small *c*. As *c* increases 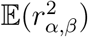 decreases. If alleles from both loci are assembled into haplotypes completely independently from the previous generation (i.e., *c* = 1, as the theoretical maximum), the expected value in this limiting case is

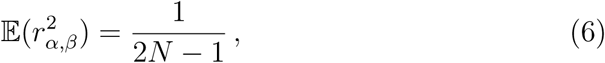

as originally derived by Haldane [9]. We state

#### Lemma 1.

*Consider a sample of n ∈*ℕ *chromosomes with two loci α, β and with fixed allele frequencies f* (*a*) = *s, f* (*A*) = *n- s at α and f* (*b*) = *u, f* (*B*) = *n -u at β. Assume that chromosomes with allele a and allele b are chosen uniformly and independently, and that the assignment of a-alleles at α is independent from b-alleles at β. Then*,

a. 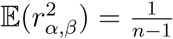
b. *This also holds if the allele frequencies s and u are randomly chosen according to discrete distributions p*_*α*_, *p*_*β*_ *on* 1,, *n* 1.
c. *In the case of fixed allele frequencies (as in* (*a*)) one has

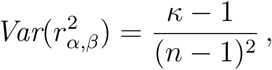

*where κ is the fourth standardized moment (kurtosis) of a hypergeometric random variable*.

*Proof*. In the scenario described in Lemma 1 (*a*), the number of individuals of type (*a, b*) is distributed as a hypergeometric *H*(*n, s, u*) random variable *X*. Thus, the expected squared correlation can be expressed by the formula

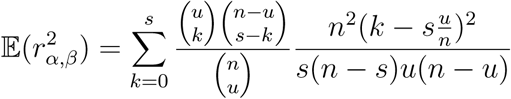

The denominator of the term 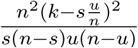 is independent of *k* and may be extracted from the summation. The remainder of the summation can then be written as

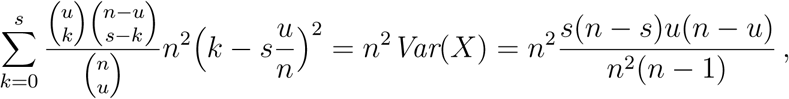

such that the initial equation can be transformed to

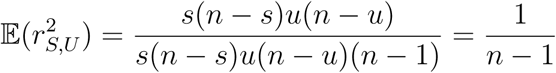

To prove Lemma 1 (*b*), it suffices to note that the same expectation with respect to the distributions of *s* and *u* can be expressed by

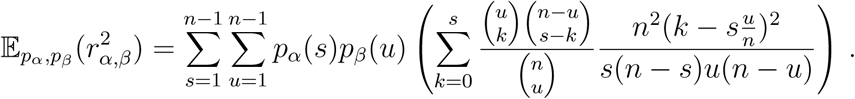

Since both distributions integrate to 1, the result remains unchanged. Finally, we obtain Lemma 1 (*c*) by first writing

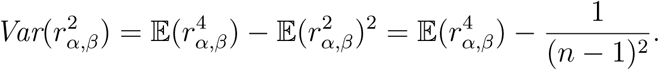

Then,

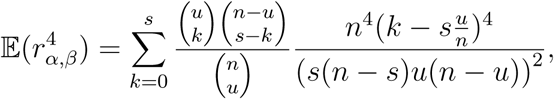

which we can identify as

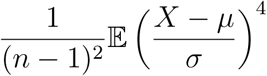

where *X* is the hypergeometric random variable mentioned above, 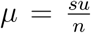 is its expectation and 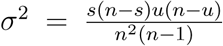 its variance. Therefore, the last expression is 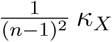, and *κ*_*X*_ is the kurtosis of *X*.

Determining 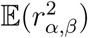 with respect to *c* for the non-limiting case is nontrivial. Sved [3] proposed an approximate solution by relating 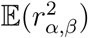 to the *conditional probability of linked identity by descent*, denoted by a parameter *Q*. He obtained the approximation

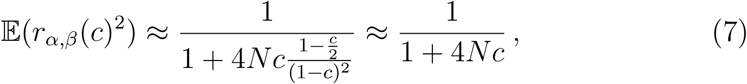

where the last approximation holds for *c <<* 1 (see also [10]).

Despite the later attempts (e.g. [11]) to improve this formula, the unsatisfactory discrepancy between eqs (6) and (7) still persists.

### 2.2. Topological Linkage

A slightly modified concept of linkage disequilibrium can be defined very intuitively in the framework of coalescent theory. Consider a sample of *n* recombining chromosomes of unit length and with recombination rate *ρ* = 2*Nc >* 0 between the ends of the chromosome. The genealogical history of such a sample can be modelled by the *ancestral recombination graph* (ARG) *A*_*n*_ [12]. *The projection of the ARG on any point γ∈* [0, 1] yields a Kingman coalescent *G*_*γ*_. Trees at different positions *γ* and *γ’* may be different due to changes of tree topology by recombination. In fact, (*G*_*γ*_)_*γ∈*[0,1]_ can be viewed as a (non-Markovian) stochastic process on the set of Kingman coalescent genealogies with state changes caused by recombination events. The genealogy *G*_*γ*_ persists for some nonempty interval [*a ≤ γ, γ ≤ b*) with expected length 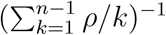. We call *S* = [*a, b*) a *segment* of the chromosome and denote its genealogy by *G*_*S*_.

As a Markovian approximation of the ARG, the so-called *Sequential Markov Coalescent* (SMC) was introduced by [13]. In this construction *A*_*n*_ as the unified history of coalescence and recombination events of the sample is omitted. Instead, genealogies change along the chromosome by uniformly choosing a branch of the current genealogy, removing the subtree below and replacing it somewhere else in the tree. This *prune-regraft* operation will be described in more detail below.

The *SMC*-construction is considered a reasonably accurate approximation of the *ARG* process, with the advantage that computation is much easier. Under the *SMC*-construction (*G*_*γ*_)_*γ∈*[0,1]_ becomes a continuous-time Markov chain, where the distance between changes in *G*_*γ*_ is the average segment length 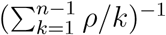, as in the *ARG*. For the remainder of this manuscript we assume that (*G*_*γ*_)_*γ∈*[0,1]_ is a realization of the *SMC*.

Any genealogy *G*_*S*_ extracted from (*G*_*γ*_)_*γ∈*[0,1]_ and valid on some segment *S* naturally provides a separation of the sample into two disjoint sets: Let *S*_1_ contain all sample members found on the “left-hand” side, and *S*_2_ all those on the “right-hand” side of the root node of *G*_*S*_. Note that “left” and “right” can also be interpreted as two different alleles, originating by a point mutation along one of the root branches. Moving from segment *S* to another segment *U* along the sequence (*G*_*γ*_)_*γ∈*[0,1]_, the genealogical tree *G*_*U*_ may differ from tree *G*_*S*_ as a result of recombination. As a consequence, also the left (*U*_1_) and right (*U*_2_) descendants below the root of *G*_*U*_ may differ from *S*_1_ and *S*_2_. Correlation can be measured in a way analogous to conventional linkage disequilibrium: Let

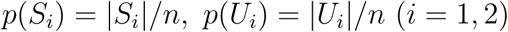

and

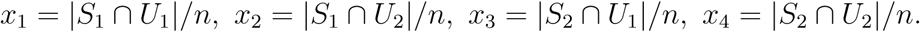

Then we call

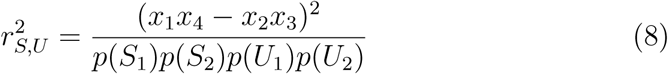

*topological linkage disequilibrium of the segments S and U*, in short *tLD*.

The choices of “left” and “right” are arbitrary, as much as the naming of alleles, and does not affect the value 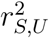. Note, that we have 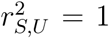, if and only if *S*_1_ = *U*_1_ or *S*_1_ = *U*_2_, where segments *S* and *U* may be different.

Such a configuration is called *complete linkage*.

*tLD* combines the concepts of *LD* and coalescent. Although *tLD* and *LD* have similar properties, they differ, for instance, in their rate of decay with distance. While any recombination event and point mutation may affect *LD, tLD* is affected only by topological changes at the root of a coalescent tree. We explore this in more detail now.

## 3. Theory

### 3.1. Limit behaviour of tLD in the SMC

Disregard for a moment branch lengths of individual coalescent trees *G*_*γ*_ of the *SMC* and consider only their branching pattern. Such trees are also called *coalescent tree topologies* (see [14] and Figure 3). The set 𝒯_*n*_ of coalescent tree topologies of *n* chromosomes has size *|*𝒯_*n*_*|*= *n*!(*n* − 1)!2^1*-n*^ [15]. Omitting branch lengths, but introducing integer-labellings of internal tree nodes which respect the time-order of these branchings, Kingman’s coalescent induces the uniform distribution on *𝒯*_*n*_. Note that the sequence of topologies, (*T*_*γ*_)_*γ∈*[0,1]_ with *T*_*γ*_ *∈𝒯*_*n*_ for *γ∈* [0, 1] that is induced by the *SMC*, is a *finite-state* Markov chain.

**Figure 1:**
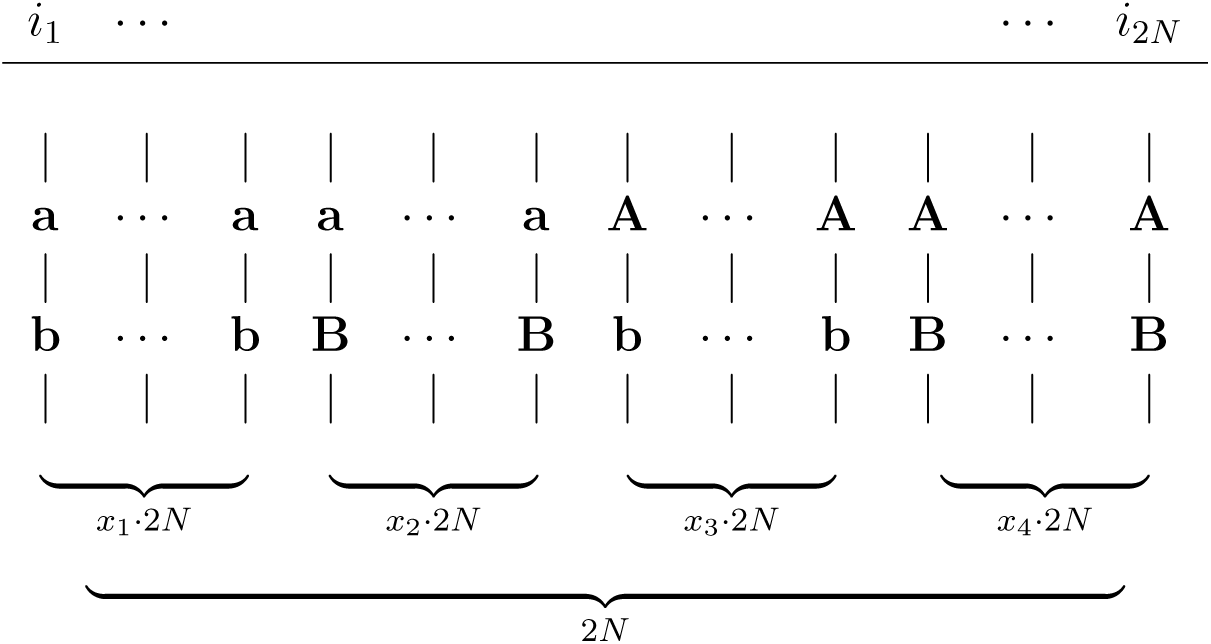
Alignment of 2*N* haplotypes *i*_1_,, *i*_2*N*_, indicated by vertical lines, with two bi-allelic loci. Frequencies given below the parentheses.

**Figure 2:**
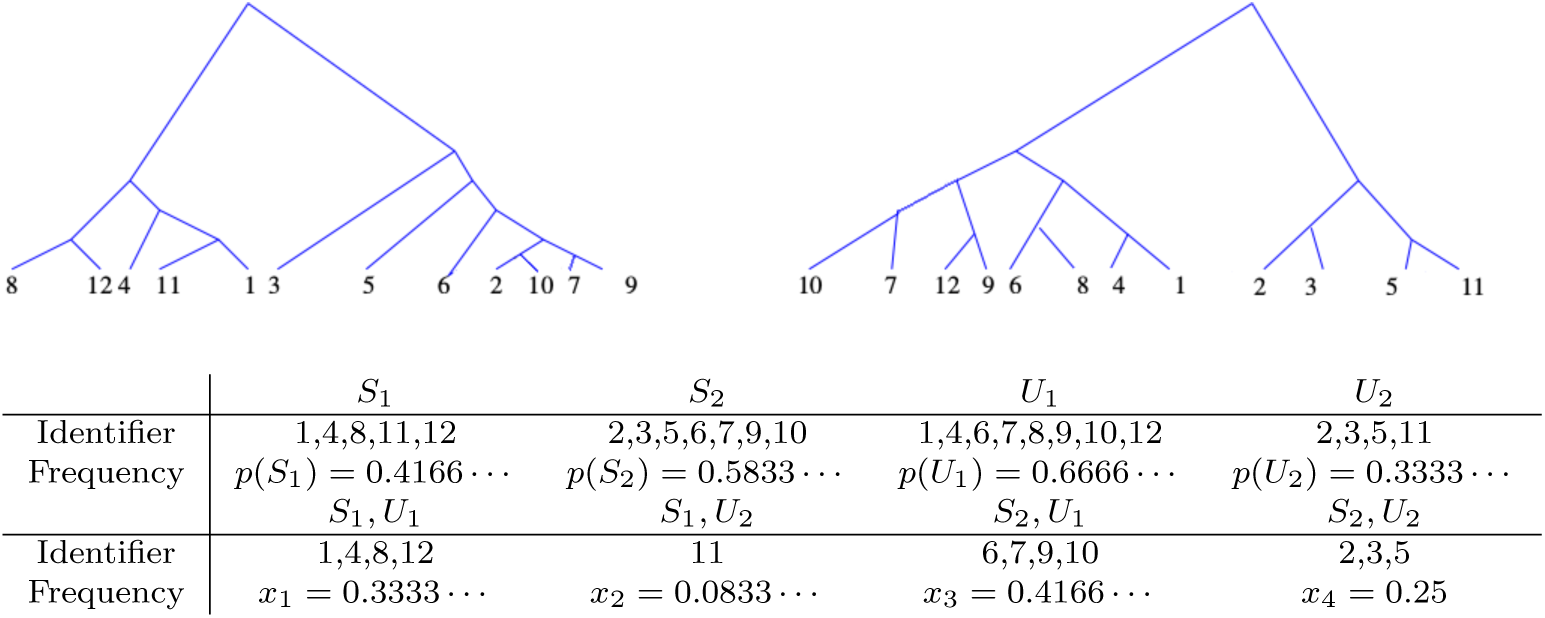
Example of the topological association of twelve chromosomes 1 − 12 in two random coalescent trees. The above configuration yields an *r*^2^-coefficient of 0.04375.

**Figure 3:**
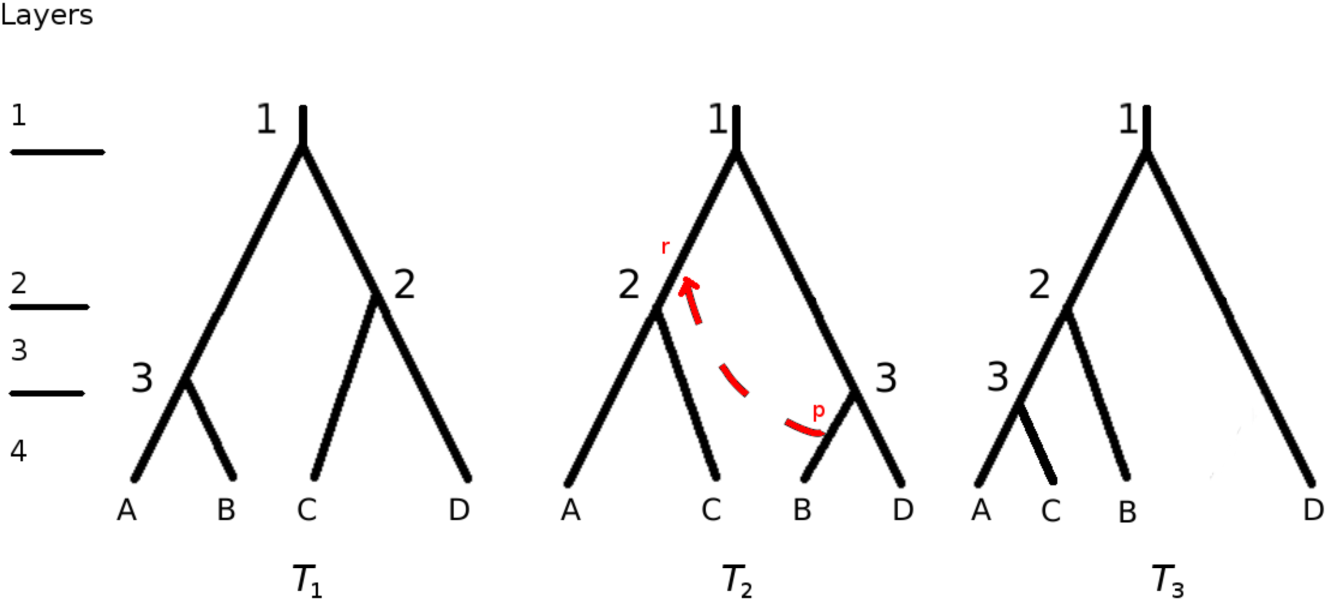
*Left:* Example of a Coalescent tree topology (*T*_1_). It is a binary tree equipped with an integer labelling representing the order of coalescent events backward in time, and identifiers (*A*-*D*) for the chromosomes at the bottom. *Middle, Right:* A *srARG* of size 4, represented by its associated topologies *T*_2_ and *T*_3_. A subtree of *T*_2_ is selected for pruning (*p*) and is regrafted (*r*) at a layer equal or smaller than where *T*_2_ was pruned. Note that the prune-regraft operation affects the internal labelling of *T*_3_.

The *k*-th *layer* of a coalescent tree topology *T* is the tree slice in which it has *k* branches (Figure 3). In particular, the 1-st layer is the (imaginary) slice with only one branch, which extends from the root back into the past. Treating each segment of a branch from the original Coalescent genealogy extending between two layers *k, k* + 1 as a seperate branch in the associated Coalescent tree topology, each topology of size *n* has exactly 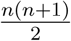 branches. Under the *SMC, T*_*γ*_ switches states at recombination sites *γ* according to a prune-regraft operation: select a branch at layer *k*, say, to place a pruning site on the tree; cut the underlying subtree; select a branch at a layer less than *k* to place a re-grafting site; re-attach the cut subtree at the regrafting site. We call such a transition a *single recombination SMC*, for short *srSMC*. Assuming an ‘infinite recombination site’ model, the complete *SMC* can be viewed as a sequence of *srSMC* s.

Consider a random *srSMC* on *n* chromosomes and extract the topologies *T*_1_ and *T*_2_ of the two coalescent genealogies *G*_1_ and *G*_2_. *T*_1_ and *T*_2_ are still uniformly distributed, but they are not independent. Given a random topology *T*_1_, assume a pruning event in layer *k* at a randomly chosen branch *b*_*k*_. Assume the branch *b*_*j*_ for regrafting to be chosen *uniformly* from the 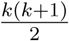 possible branches. A *srSMC* can be viewed as a triplet (*T*_1_, *b*_*k*_, *b*_*j*_). To obtain the conditional distribution *P* (*T*_2_ *T*_1_ = *T*) one has to average over all possible pruning and re-grafting sites. The probability of pruning in layer *k >* 1 is given by (see [14])

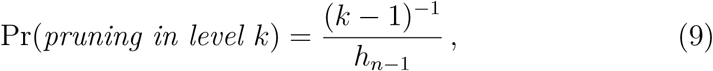

where 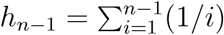 is the (*n* − 1)-st harmonic number, keeping in mind that this probability does depend on the duration of the layer, i.e. on the coalescent branch lengths. The prune-regraft-operation describes how states in the Markov chain (*T*_*γ*_)_*γ∈*[0,1]_ are changed. In particular, the set of all *Aldous moves* on a topology *T*_*γ*_ (see [16]) is a subset of the possible transformations *T*_*γ*_ can undergo. It follows that (*T*_*γ*_)_*γ∈*[0,1]_ is recurrent and aperiodic. We now consider the leftmost and rightmost chromosomal segments *S* = [0, *γ*_1_] and *U* = [*γ*_2_, 1], *γ*_1_ *≤ γ*_2_, along the interval [0, 1]. We have the following result on the limiting behaviour of the squared correlation 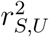.

#### Theorem 1.

*Given segments S and U with topologies T*_*S*_ *and T*_*U*_ *and the topological groupings* (*S*_1_, *S*_2_) *and* (*U*_1_, *U*_2_). *Then*

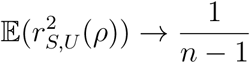

*for ρ → ∞*.

*Proof*. (*T*_*γ*_)_*γ∈*[0,1]_ is a Markov chain with stationary distribution 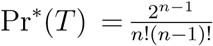 for all coalescent tree topologies *T*. Since it is recurrent, there exists an integer *M ∈*ℕ (*mixing time*), depending only on the sample size *n*, such that for *ϵ >* 0

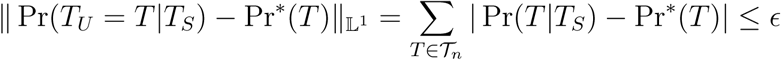

given there are *M* or more state changes in (*T*_*γ*_)_*γ∈*[0,1]_. By choosing *ρ* sufficiently large, it is possible to ensure that the probability of *M* or more changes is close to 1. Therefore, Pr(*T*_*U*_ = *T|T*_*S*_) may be brought arbitrarily close to Pr*\**(*T*).

Let *ρ → ∞* and consider the random variable *k*_*ρ*_ = *|S*_1_ *∩U*_1_*|*of individuals that are on the left of both trees *T*_*S*_ and *T*_*U*_ under the *SMC* with recombination rate *ρ*. As *ρ→∞*, *k*_*ρ*_ converges in distribution to a random variable *X* which, given *|S*_1_*|* and *|U*_1_*|*, is distributed 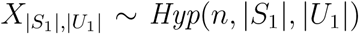. *|S*_1_*|* and *|U*_1_*|* themselves are uniformly distributed on {1, …, *n -*1}.

Therefore, Lemma 1 is applicable to the limiting r.v. *X*. Convergence in distribution of the *k*_*ρ*_ implies convergence of 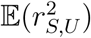 to 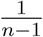.

Over large distances, we therefore recover the limiting value predicted by Haldane [9], see eq (6). Note that by item (*c*) of Lemma 1 we can also obtain an exact expression of the variance of *tLD* in this situation.

### 3.2. Decline of tlD with distance

Regarding the expected decline of *tLD* along the chromosome, we state the following

#### Lemma 2.

*Let T*_1_, *T*_2_ *be the two topologies resulting from a srSMC. The probability of a topological change, i.e. breaking of complete linkage, between T*_1_ *and T*_2_ *is asymptotically* 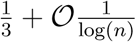.

*Proof*. Recombination events that have an effect on *tLD* can be subdivided into two groups: events which shift a non-root branch above the root or events which move a branch from the left to the right (or vice versa) root-subtree without changing the root. We call the latter *switching events*. The probability of root-changing events is 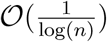 [14]. To calculate the probability Pr(*switch*) assume w.l.o.g. that a branch is moved from left to right. Suppose pruning takes place in layer *k*. The right side has 1 *≤r≤ k -*1 branches, where each number *r* has probability 1*/*(*k -*1). The probability of selecting a branch on the left for pruning is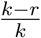, and some arithmetics leads to the probability of selecting a branch on the right for regrafting, which is 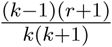 when averaged over all *k*-sized coalescent tree topologies. This needs to be multiplied by 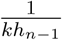 (compare equation 9), and then summed over all levels *k*. Therefore, we obtain

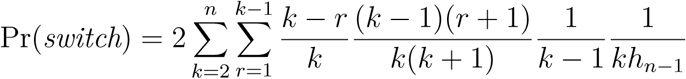

where the factor 2 accounts for the two possibilities, switching from left to right or vice versa. After some simplifications, this can be rewritten as

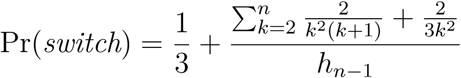

The series in the numerator converges to (4*π*^2^ –33)*/*9 *<* 1 and *h*_*n*_*∼* log(*n*), which conclude the proof.

Knowing the proportion of recombination events contributing to the decay of 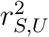, we calculate now the expected proportion of chromosomes affected by a switching event between two segments. Denote by *L*_*S,U*_ the number of chromosomes whose assignment to either the left or right class has *not* been affected by switching. Note that the quantity *L*_*S,U*_ is similar to *Q* in [3]. However, we are here interested in a *joint* probability of identity by descent, and not in a probability conditioned on identity at one of the loci. *L*_*S,U*_ = *n* means perfect linkage and *L*_*S,U*_ *< n* means some switching took place (Figure 4). Lemma 1 is not readily applicable here, because its conditions require that *L*_*S,U*_ vanishes.

**Figure 4:**
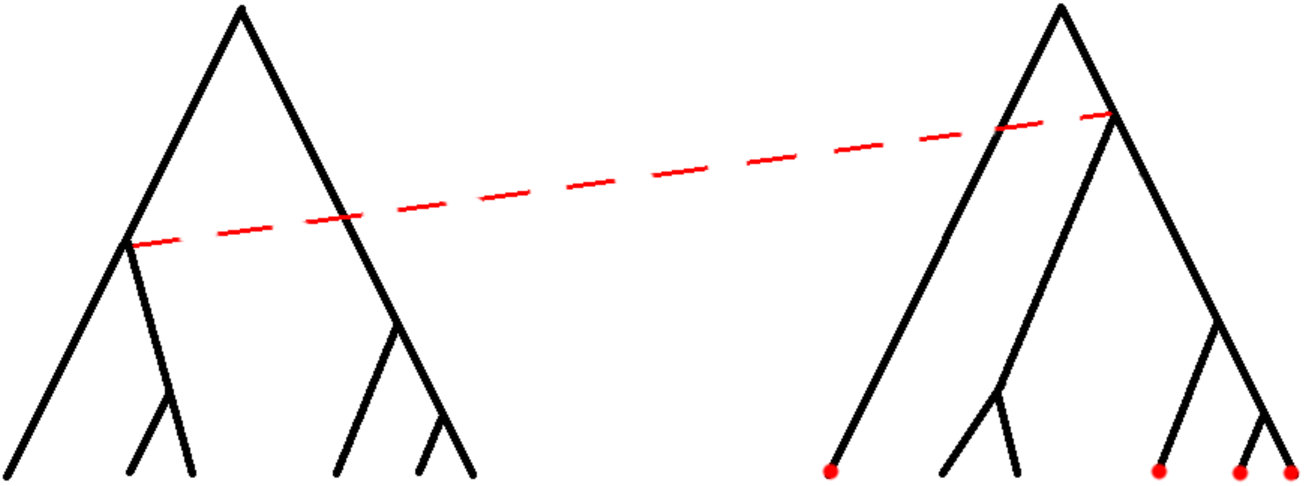
A single recombination event moves a branch of the left root-subtree to the right side. In the resulting tree, chromosomes marked by red dots remain in the same left/right-grouping as before recombination took place. Their number is *L*_*S,U*_.

#### Lemma 3.

**𝔼** (*L*_*S,U*_) *declines geometrically with rate approximately* 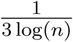 *per recombination event*.

*Proof*. According to Lemma 2 about 1*/*3 of all recombination events move some individuals from one side of the tree to the other. To determine how many are moved, it suffices to note that recombination events are distributed uniformly over the branches of a given genealogy *T* in the *SMC*, such that the size of the subtree *T*_*r*_ below a recombination event is distributed according to the neutral frequency spectrum [17]. Thus,

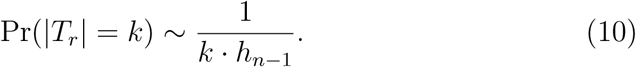

The expected proportion of chromosomes affected by a recombination event is the expectation of the above distribution divided by *n*, i.e.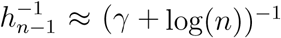, where *γ* is Euler’s constant. We can combine those two results in a recursion formula for 𝔼 (*L*_*S,U*_). Let *U*_1_, *U*_2_ be two neighboring segments, then we have

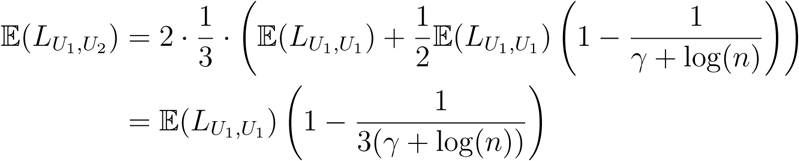

Iterating this formula with initial value

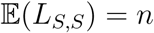

shows that

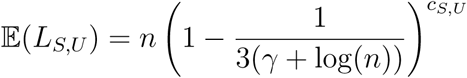

with *c*_*S,U*_ representing the number of recombination events encountered moving along the chromosome from *S* to *U*, which depends only on *ρ*.

### 3.3. Numerical approximation of 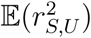

The above results about the decline of the parameter *L*_*S,U*_ suggest an approximation scheme for the expectation of *tLD* with respect to the number of recombination events seperating two segments. *L*_*S,U*_ can be written as *L*_*S,U*_ = *l* + *m*, where *l* (*m*, respectively) is the number of *L*_*S,U*_-chromosomes on the left (right) side of both trees. There are *p* = *|S*_1_*|–l* additional individuals on the left side of *T*_*S*_ and *q* = *|U*_1_*|–l* on the left side of *T*_*U*_. See Figure 5 for an example.

**Figure 5:**
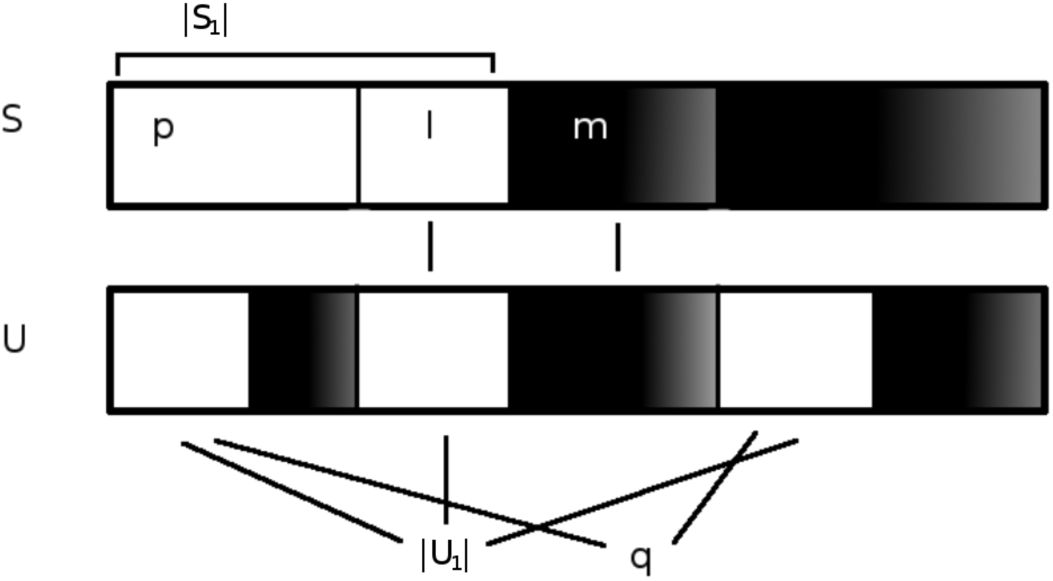
At segment *S*, the sample members are grouped by tree topology into left (*white*) and right (*black*). Moving from *S* to *U*, a number of *L*_*S,U*_ = *l* + *m* members remains unaffected by recombination, while the rest of the members might switch sides.

To calculate 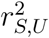, we need to determine how many of those additional *p* chromosomes on the left side of *T*_*S*_ are also on the left side of *T*_*U*_ by chance. We assume that this number is given by a hypergeometric distribution. This is an approximation, because in both *ARG* and *SMC* the number of possible left-right configurations is smaller than the number of all hypergeometrically possible configurations. However, this restriction becomes negligible with increasing distance between *S* and *U*, i.e. with increasing number of recombination events. Let *k* denote the number of individuals which end up on the left side of *T*_*S*_ and *T*_*U*_ by this hypergeometric assignment, such that there are *k* + *l* chromosomes in total that are on the left side of both trees.

Under these assumptions, the expected *tLD* is

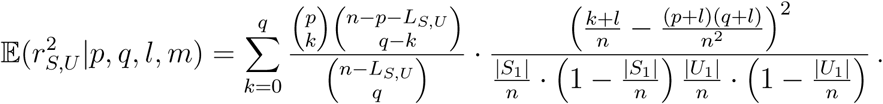

It is useful to note that the term

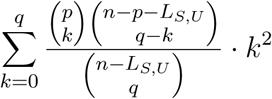

is the second moment of a hypergeometric variable with parameters *n– L*_*S,U*_, *p, q* and with expectation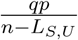. We calculate therefore

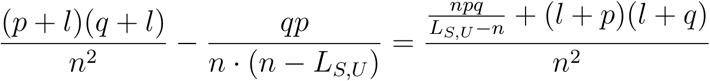

and use this to rewrite the original expression as

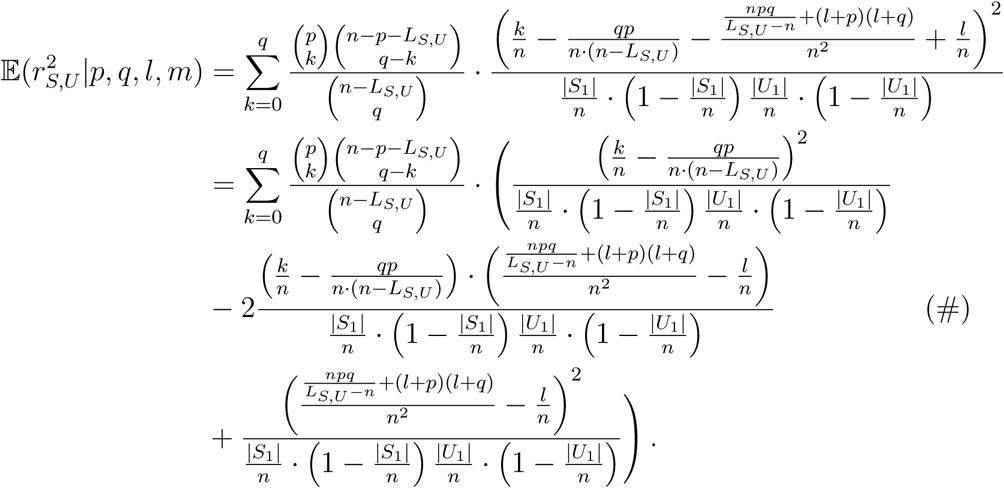

This expression can be simplified. The middle term of the summation (line # above) vanishes because of symmetry; the first summand contains the variance of a *Hyp*(*n–l–m, p, q*) random variable divided by some constant terms. Thus, we arrive at

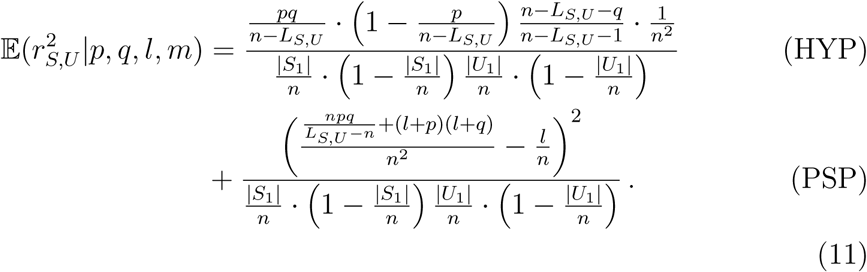

In this form, the contribution that arises from the hypergeometric random assignment, labelled HYP, and the remaining parameter-specific (PSP) terms are separated. This decomposition is useful in two ways. First, under the above assumptions, an upper bound can be obtained for 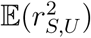 at least if *L*_*S,U*_ is small in relation to *n* (See Lemma 4). Second, by averaging over all configurations it is possible to approximate 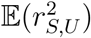 regardless of the tree topologies at segments *S* and *U*. Since the size *|S*_1_*|* of the left side of *T*_*S*_ is uniform on {1,, *n –*1}, we start by choosing *|S*_1_*|* randomly according to the uniform distribution. The *L*_*S,U*_-sized portion of individuals not having undergone recombination when going from *S* to *U* is then subdivided into *l* individuals which are on the left side both in *T*_*S*_ and *T*_*U*_, and *m* individuals which are on the right side in both trees by choosing hypergeometrically from the assignment at *S*, which implicitly determines the parameters *p, l* and *m*. The number *q* of additional individuals on the left side of *T*_*U*_ is determined by drawing uniformly from {1, …, *n L*_*S,U*_}. These calculations are easily performed by computational algebra. Note that the explicit calculation of SNP-based *LD* is much harder because the sizes of the classes are not uniform. The resulting approximation of the expected *tLD* with respect to *L*_*S,U*_ has to be scaled with respect to the expected decay of *L*_*S,U*_ itself. Assuming that recombination events are uniformly distributed across a chromosome, the approximation of 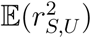 can be expressed in terms of physical distance in base pairs, given a constant recombination rate per bp.

### 3.4. An upper bound to 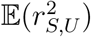

#### Lemma 4.

*Assume that the parameter L*_*S,U*_ *is small in relation to the sample size n. Then, we have:*

a. *The expectation of the parameter-specific contribution (“PSP”) in equation 11 is of order* 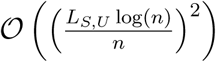.
b. *The proposed approximation of* 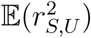 *is bounded from above by* 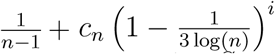, *where i denotes the number of recombination events seperating S and U and c*_*n*_ *is some constant*.

*Proof*. Under the assumption of *L*_*S,U*_ *< n*, it is possible to write 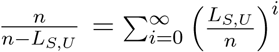. Furthermore, *l ·* (*n - p - q - l*) *∈* [*-l · n, l · n*]. This allows us to rewrite the numerator of the parameter-specific term in the following way:

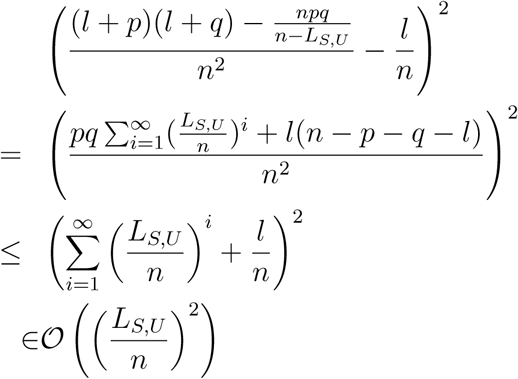

The last statement is true because *l/n ≤L*_*S,U*_ */n*.

Under the assumptions from section 3.3, and if *L*_*S,U*_ is small in comparison to *n*, then the sizes *|S*_1_*|* and *|U*_1_*|* of the left sides of the genealogies are approximately independent and uniformly distributed on {1, …, *n –*1}. Thus, the expectation of the denominator in

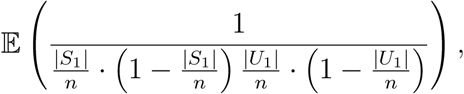

converges to

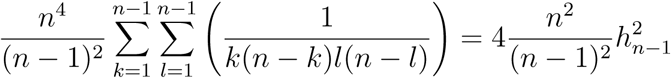

as *L*_*S,U*_ */n* becomes small. This term is of order 𝒪(log(*n*)^2^), allowing us to conclude that

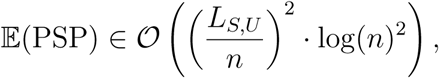

establishing claim (a).

To show (b), we recall Hölder’s inequality

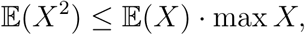

for a nonnegative random variable *X*. Let 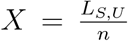, assuming exactly *i* recombination events between *S* and *U*. The maximal value of this random variable is 1 (no decline at all). The expectation of *X* given the number of recombination events *i* between *S* and *U* has been calculated in section 3.2 as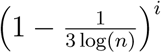. Thus, 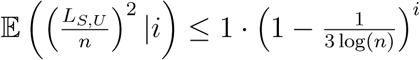, and therefore

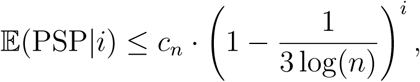

with *c*_*n*_ *∈*ℝ depending on *n* and of maximally (squared) logarithmic growth. The expectation of the hypergeometric contribution is 0 for *L*_*S,U*_ = *n* and converges to 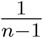 from below for *L*_*S,U*_ */n →* 0, finishing the proof.

The above calculations hold in the setting assumed in 3.3 and are an approximation of the true *SMC*. Furthermore, since the term 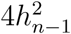 is not bounded from above, this upper bound is only of relevance for large *n*.

## 4. Simulations and Application

We use the program ms [18] to generate samples of *n* chromosomes with recombination rate *r* and mutation rate *θ*. Computing *tLD* requires two steps. First, select two disjoint loci (sub-intervals of [0, 1]). Second, determine the top-most left and right root sub-tree clusters for both loci. We consider two possibilities: (a) use the true tree structure provided by the coalescent simulations (option “-T” selected) or (b) estimate the clusters from SNP data. In the latter case, this is achieved by a two-means clustering approach: We first determine the two most diverged haplotypes (‘antipodes’) and then assign the remaining haplotypes to either of the antipodes based on minimal Hamming distance. The estimated clusters agree well with the true (simulated) clusters if sufficient SNPs are available for estimation. A minimum number of 10 SNPs gives good results (M. Rauscher, unpubl. data; see Figure 7). This is confirmed by the excellent agreement of the summary statistics (average and variance) of *tLD* determined from actual and estimated clusters (see Figure 6) and by the very good agreement of individual *LD* estimates (see Figure heatmap). Hence, true and estimated values of *tLD* agree in general. Importantly, this observation does not depend much on distance. However, the correlations depend heavily on the relation between mutation and recombination rates.

**Figure 6:**
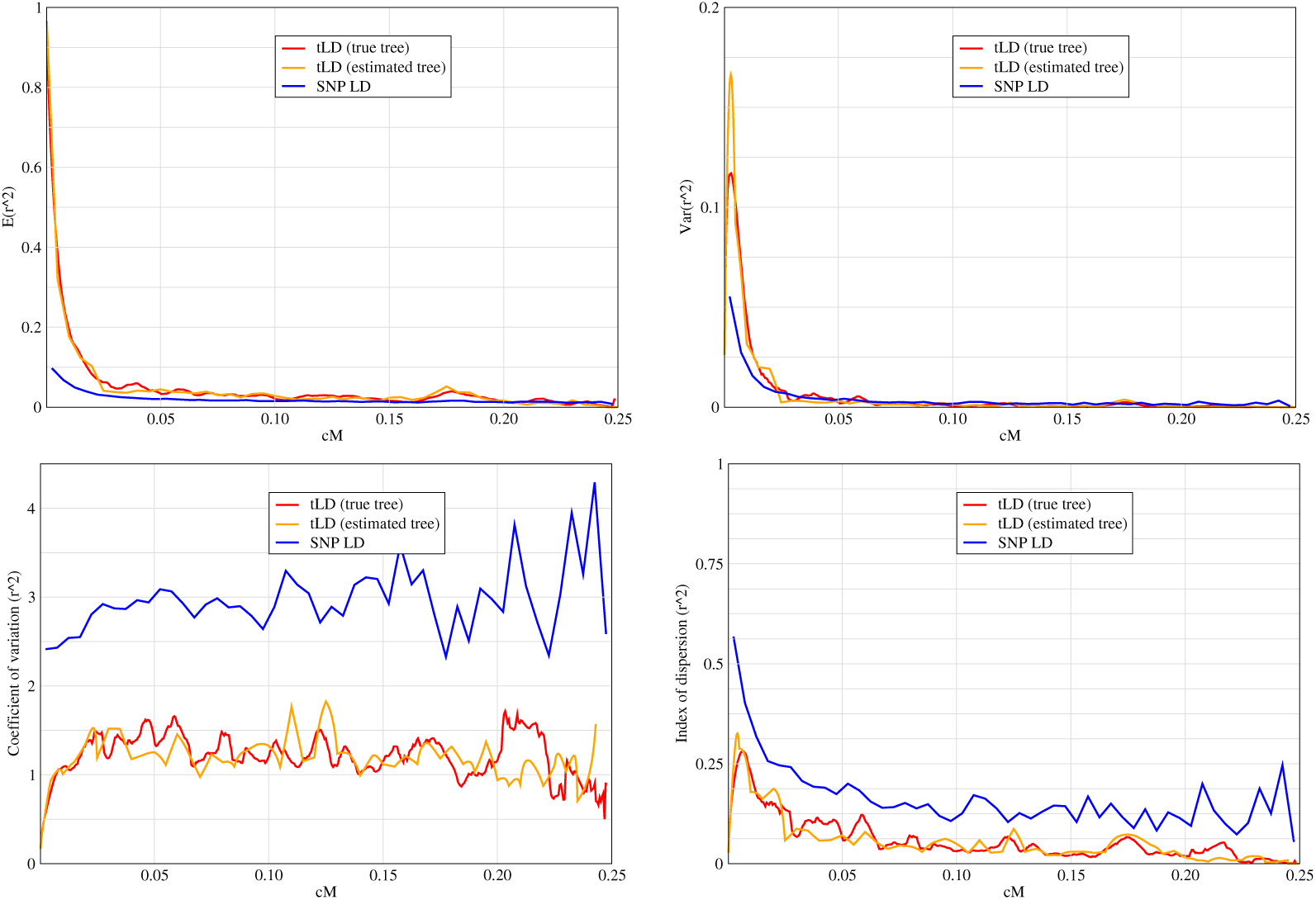
*tLD vs*. SNP-*LD*. Average (top left), variance (top right), coefficient of variation (bottom left) and index of dispersion (bottom right) of *r*^2^. Data from a single simulation run performed with the program ms [18]. Parameter settings: ms 200 1 -t 100 -r 100 1000 -T. For a poulation size of *N* = 10^4^ and a recombination rate of 1*cM/M b* the simulated region corresponds to 0.25cM or 250kb physical distance. Red: *tLD* calculated from the actual coalescent trees (i.e., using the trees obtained by setting the parameter *-T*). Orange: *tLD* calculated from estimated tree topology (see text). Blue: Conventional *LD* calculated from SNP pairs. Coefficient of variation: *σ/µ*; index of dispersion *σ*^2^*/µ*.

**Figure 7:**
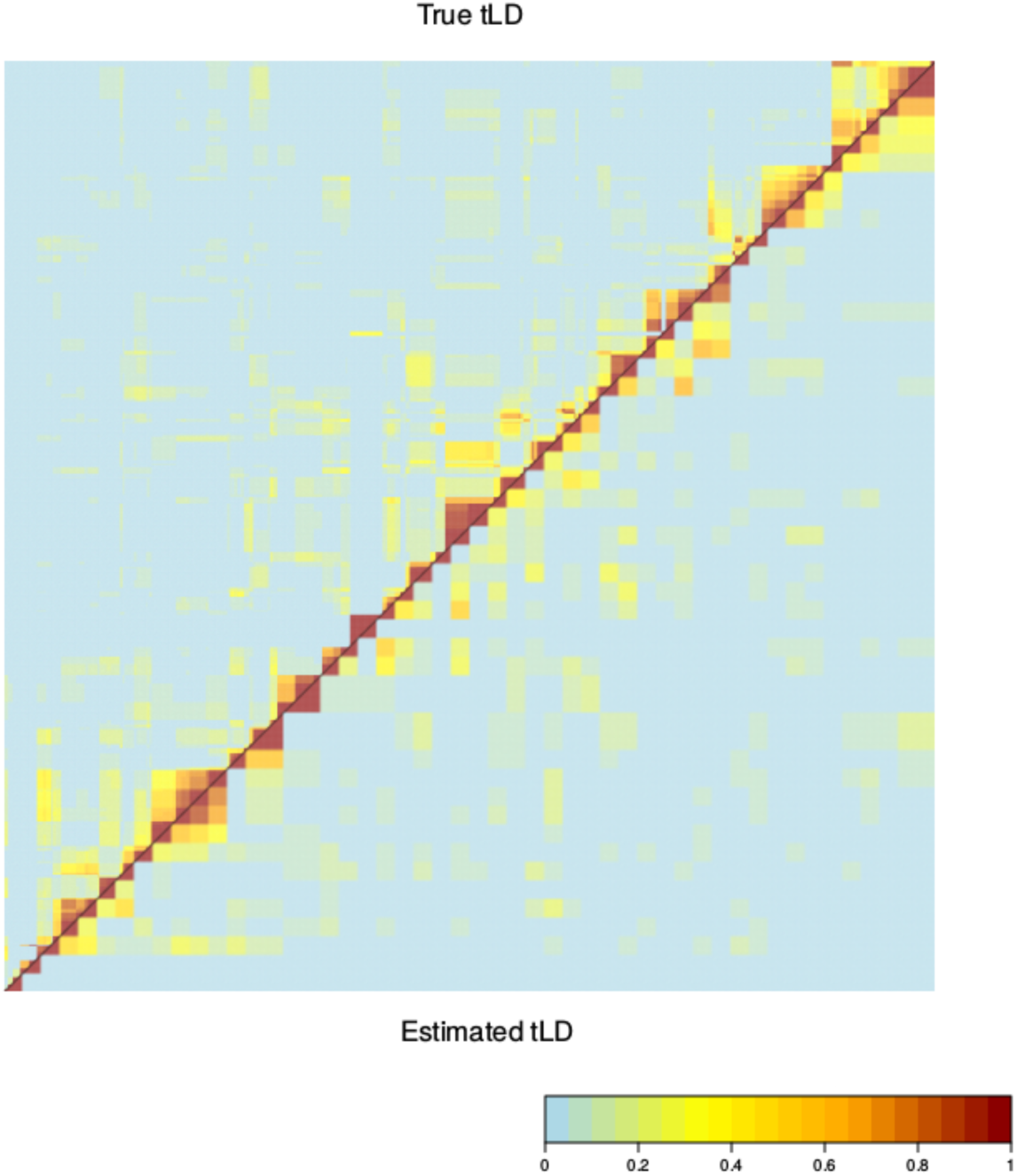
Heatmaps of true *tLD* calculated from tree topologies and *tLD* calculated from estimated tree topologies, performed on the same simulated dataset used in Figure 6

Comparing *tLD* with conventional *LD* in simulated data, we find that *tLD* declines comparably slowly and smoothly, although the variance is large. In contrast, conventional *LD* vanishes very quickly over similar distances (Figure 6). Although average *LD* is small, its variance is high, in particular for short distances. In this regime the variance of *tLD* is much smaller relative to the average than that of conventional *LD* (coefficient of variation *σ/µ* shown in Figure 6). These observations are theoretically supported by Lemma 2.

In practice tree topology is unknown. To estimate tree topology at the tree root, we apply the two-means clustering approach described above. We use the human 1*k* genomes data [19] to determine *tLD* across the LCT region on chromosome 2 in the CEU (Central Europe) and YRI (Yoruba) populations. We estimate *tLD* using a window size of 5*kb* per locus. Most of these windows contain 10 or more SNPs. We find a strongly elevated level of *tLD* in the CEU population compared to YRI. As can be seen from the heat maps, there is a much higher, and a longer-ranging level of correlation to be observed for *tLD* than for conventional *LD* (Figure 8).

**Figure 8:**
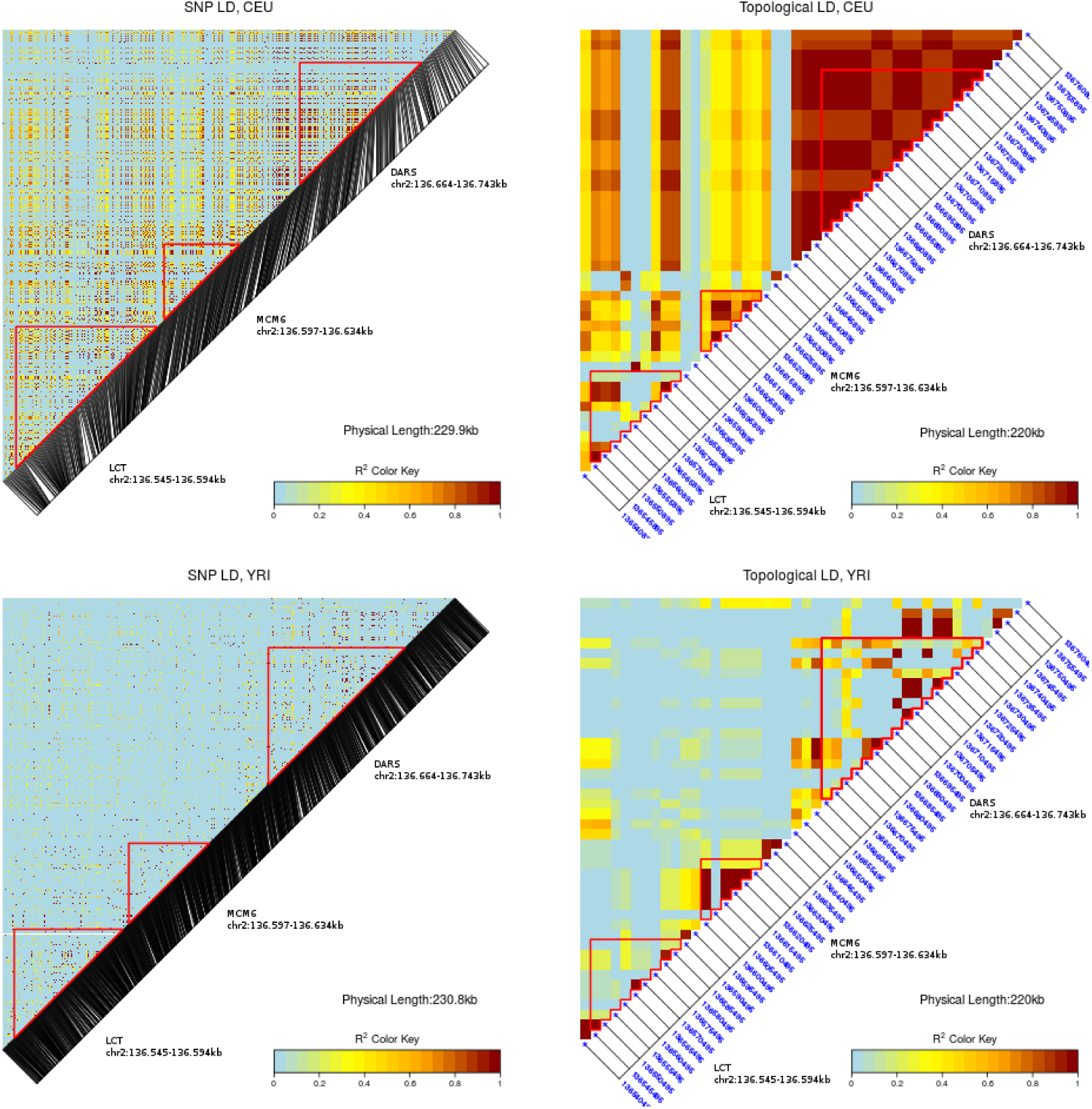
The heatmaps of conventional (left) and topological *LD* generated for the LCT and neighbouring regions of the CEU (top) and YRI subsamples of the human 1k genomes project [19].

**Figure 9:**
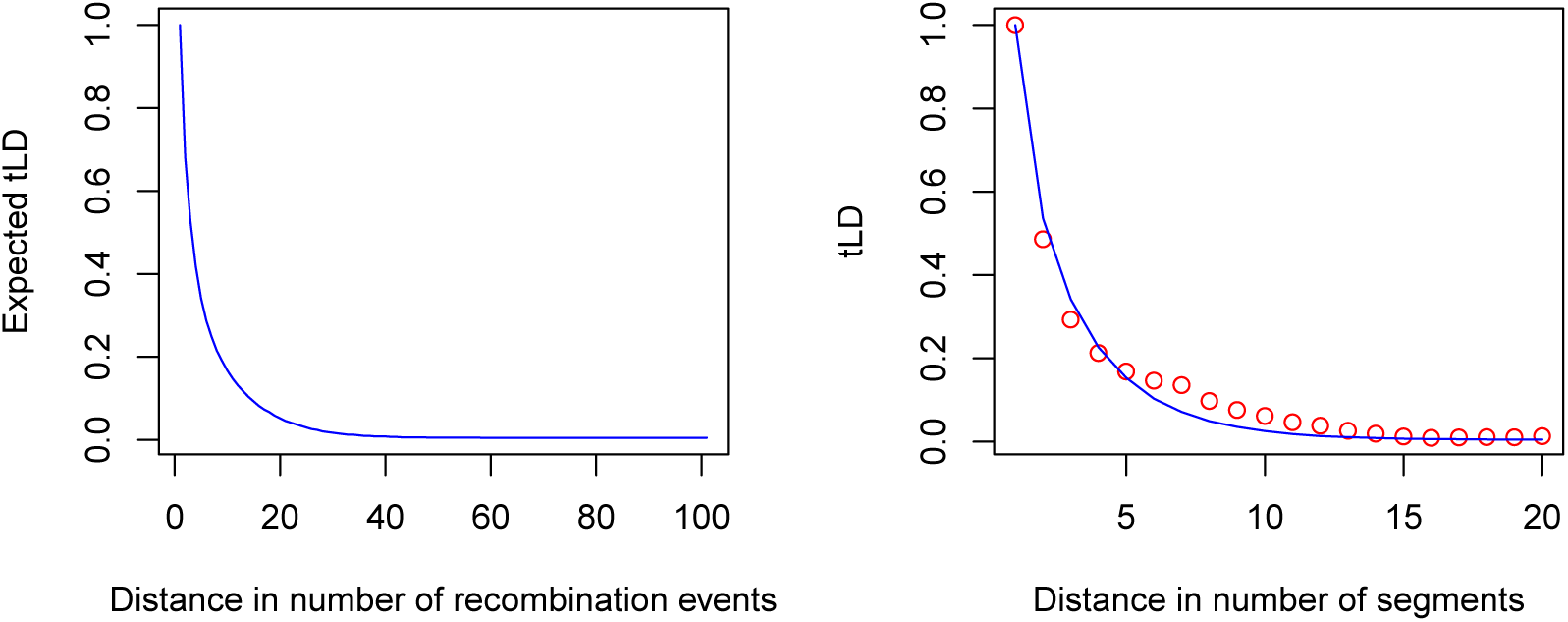
*Left:* 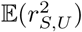 in a sample of size *n* = 100 with respect to *L*_*S,U*_ declining at rate 1*/*(3 log(100)) with respect to the number of recombination events encountered in the sample. *Right, Dots:* Average *tLD* between pairs of segments with respect to their distance in units of 2500*bp* (segment size: 5000*bp*), estimated on the region expanding over approx. 5.51 *·* 10^6^ *-* 5.52 *·* 10^6^ on chromosome 5 of the human 1k genomes data (CEU). *Blue Line:* Least-squares fit of expected *tLD*, estimating 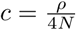 at *≈* 1.116*cM/M b*.

## 5. Discussion

We have introduced the concept of topological linkage disequilibrium (*tLD*). Like conventional *LD* it is a means to analyse and quantify chromosomal linkage. In contrast to the latter *tLD* can in principle be calculated for arbitrary pairs of loci, even if not polymorphic, because the concept hinges on coalescent tree topology, or subtree clusters, not on allele pairs. In practice, however, tree topology is usually not directly observable and needs to be estimated from polymorphism data. A moderate number of SNPs, on the order of ten, turns out to be sufficient to determine the two root-subtree clusters (assuming a binary coalescent tree) quite reliably (Figure 7). Given SNP data, we use a 2-means clustering approach, described in more detail above. Even with a single SNP, the correct root-subtree clusters may be recovered. In fact, this happens with probability roughly equal to (1 + 1*/*2)*/h*_*n–*1_, i.e. with the probability that a SNP lands on either one of the two root branches. This is about 1*/*2 for *n* = 12 and about 1*/*4 for *n* = 227. Alternatively, one may also use other kinds of markers, for instance microsatellites [20], or structural variants, if available, to estimate the root-subtree clusters.

As a function of distance between loci (or markers) average *tLD* and conventional *LD* behave qualitatively similarly. Both decay toward the same limit, as expressed in Haldane’s equation (see eq 6). However, *tLD* decays more slowly, and is less dispersed, than conventional *LD* (Figure 6). The coefficient of variation of *tLD* is only slightly larger than one, almost independently of distance between loci, and only about a third of that of conventional *LD*. Conversely, the inverse of the coefficient of variation, in signal processing also called ‘signal to noise ratio’, is about three times as high for *tLD* than for conventional *LD*.

The concept of *tLD* is embedded in coalescent theory. The first moment of *tLD* and its limiting behavior can be analytically approximated by a simple function using arguments derived from (Kingman-)coalescent properties. A currently open problem is to integrate the concept of *tLD* into ancestral recombination graph (*ARG*) theory. Here, we have resorted to its Markovian approximation, represented by the *SMC* to derive the presented results. We hypothesize that similar results must hold for the *ARG* in terms of limiting behaviour. In the ARG setting, the decay of *tLD* is possibly somewhat slower than in the SMC setting, because in the *ARG* genealogies can be reverted back with a higher probability than in the *SMC*.

Given that the various multi-genome resequencing projects furnish now whole-genome sequences with sample sizes of several hundred or more, the concept of *tLD* should prove useful for any analysis of linkage disequilibrium and gene interactions in such experimental studies.

## Acknowledgments

This work was supported by the German Research Foundation (DFG-SPP1590).

